# Mechanistic Link Between Glyoxalase 1 Expression and Methylglyoxal-Induced Oncogenic Stress in Prostate Cancer

**DOI:** 10.64898/2025.12.05.691672

**Authors:** Alana Battad, Francisco Nunez, Mya S. Walker, Sydney Hodge, Kelani Sun, Elise Desjarlais, Edwin DJ Lopez Gonzalez, Seigmund W.T. Lai, Leanne Woods-Burnham, Stanley Hooker, Rayyan Aburajab, Qianhua Feng, Min Talley, Sharon Baumel-Alterzon, David W. Craig, Rick A. Kittles, Yun Rose Li, John Termini, Sarah C. Shuck

**Affiliations:** Departments of Diabetes and Cancer Metabolism, Beckman Research Institute at City of Hope; Departments of Population Sciences and Division of Health Equity, Beckman Research Institute at City of Hope; Departments of Integrative Translational Sciences, Beckman Research Institute at City of Hope; Departments of Radiation Oncology, Beckman Research Institute at City of Hope; Departments of Cancer Genetics and Epigenetics, Beckman Research Institute at City of Hope; Departments of Biostatistics and Mathematical Oncology, Beckman Research Institute at City of Hope; Departments of Molecular and Cellular Endocrinology, Beckman Research Institute at City of Hope; Departments of Cancer Biology and Molecular Medicine, Beckman Research Institute at City of Hope

## Abstract

Prostate cancer (PCa) is the second leading cause of cancer-related death in American men, with African American/Black (AA/B) men experiencing higher incidence and mortality than European American (EA) men. Obesity, which disproportionately affects AA/B men, is linked to increased PCa mortality, potentially through metabolic dysregulation. We hypothesize that methylglyoxal (MG), a reactive byproduct of glucose, lipid, and protein metabolism that is elevated in obesity, contributes to PCa progression. MG forms covalent adducts on DNA, RNA, and protein. We found that MG-adducts are elevated in AA/B men with PCa compared to EA men with PCa, as well as men without PCa. AA/B men with PCa had a higher frequency of SNP rs1049346 in glyoxalase 1 (GLO1), the primary MG detoxification enzyme. PCa cell lines from EA (C4-2) and AA/B (MDA-PCa-2b) men showed differential rs1049346 status, with C4-2 cells heterozygous and MDA-PCa-2b cells homozygous for the variant. This was associated with altered GLO1 expression and activity, with MDA-PCa-2b cells exhibiting reduced GLO1 function and increased MG-adducts compared to C4-2 cells. MG altered DNA repair and RNA processing pathways and induced distinct metabolic shifts in MDA-PCa-2b compared to C4-2 cells, including increased glycolysis and reduced oxidative phosphorylation. Transcriptomic analysis revealed unique MG-induced stress responses including a tenfold higher induction of TXNIP in MDA-PCa-2b vs. C4-2 cells, a gene inversely linked to GLO1 expression and activity. These findings suggest that MG stress may contribute to PCa progression in AA/B men through metabolic reprogramming and impaired detoxification, offering insight into potential precision medicine applications.

**Statement of Significance:** Patients with obesity and diabetes have an elevated risk of cancer mortality. Defining how metabolic alterations contribute to this link is critical to understanding disease progression and identifying strategies to improve outcomes.

## Introduction

Prostate cancer (PCa) is the second leading cause of cancer-related death among American men and is notably more aggressive and lethal in certain racial groups. One group particularly susceptible to this discrepancy is African American/Black (AA/B) men, who have a two-fold higher risk of dying from PCa compared to European American (EA) men.^1^ Social, systemic, and biological factors likely contribute to the mechanisms underlying this disparity. A potential confounder is diabetes, which is more prevalent in AA/B men than EA men and significantly increases the risk of PCa mortality.^2^ Diabetes may exacerbate the metabolic dysfunction that is a hallmark of many cancers, but the mechanisms linking these malignancies are not clear. A potential mechanism is the formation of covalent macromolecular adducts induced by reactive metabolic byproducts. One of the most abundant metabolic byproducts is methylglyoxal (MG), which modifies DNA, RNA, and protein to form advanced glycation end products (AGEs).^3^ The primary MG-induced adducts on DNA and RNA are *N*^2^-carboxyethyl deoxyguanosine (CEdG) and *N^2^*-carboxyethyl guanosine (CEG), respectively.^4, 5^ These MG-induced AGEs can cause DNA mutations, reduce RNA stability and translation, and alter protein stability and function.^3^ To reduce these impacts, cells utilize the glyoxalase pathway, with the rate limiting step of this pathway performed by glyoxalase 1 (GLO1).^6^ The relationship between GLO1 and PCa progression is complicated by reports of increased protein levels associated with poor prognosis and decreased transcript levels in metastatic castration resistant PCa.^7, 8^

A key limitation of prior studies is that they do not capture GLO1 enzymatic activity, which can be impacted by single nucleotide polymorphisms (SNPs).^9, 10^ In patients with type 2 diabetes, SNP rs1049346 was significantly associated with diabetic nephropathy and retinopathy, and lower GLO1 expression and activity.^11^ However, the association between SNP rs1049346 and PCa has not been addressed. To prevent the impact of increased MG-induced AGEs that result with reduced GLO1, cells utilize the soluble receptor for AGEs (sRAGE). By sequestering AGEs, sRAGE prevents AGE activation of the full-length cell surface RAGE receptor, which is a driver of cancer onset and progression.^12^

The impact of increased MG-adduct formation with reduced GLO1 can be exacerbated by cellular stress and changes in DNA repair, RNA regulation, and gene and protein expression.^13–15^ This can ultimately result in the formation of long-term genomic mutations and changes in RNA stability and translation.

Early detection and targeted treatment are crucial to combat PCa lethality, particularly in high-risk populations.^16^ Here, we present evidence that PCa in AA/B men is associated with GLO1 SNP rs1049346, a variant located in the 5’ UTR region of the GLO1 gene and found to be homozygous in the MDA-PCa-2b cell line. This variant may impair GLO1 transcription and enzymatic activity, leading to MG accumulation and cellular stress, thereby contributing to the aggressive nature of PCa in AA/B men. Here, we comprehensively characterize the relationship between GLO1 SNP rs1049346 on MG-adduct burden and GLO1 expression and activity in both patients and in cellular models.

## Materials and Methods

### Clinical Study Design

Between 2012 and 2014, men with histologically confirmed PCa and healthy age- and ethnicity-matched controls were recruited and consented to an IRB-approved study at several clinical sites in the Chicago area by Dr. Rick Kittles’ team at Jesse Brown Department Of Veterans Affairs Medical Center (JBVAMC), John H. Stroger, Jr. Hospital of Cook County (JHS), and Northwestern Memorial Hospital (NMH). From this cohort, we designed a nested case-control study including AA/B men with (n = 83) and without (n = 83) PCa and EA men with (n = 77) and without (n = 128) PCa. Demographic and clinical data were collected using questionnaires and are presented in Supplemental Table 1. All analyses were performed by blinded lab personnel.

### Experimental Treatments

Unless otherwise indicated, C4-2 and MDA-PCa-2b cells were cultured overnight in 7 mM glucose (basal condition) or 25 mM glucose (mimicking hyperglycemic condition) for 24 hours or treated with 500 µM MG for 3 hours, for a total of three treatment conditions. MG was synthesized and quantified as previously described^4^.

### Statistical analysis

The data set contains 315 subjects including 148 patients with prostate cancers and 166 subjects as controls; 1 subject has missing data on cancer status. The proportion of AA/B is 44.8% (n=141) and EA 55.2% (n=174). R Software, R-project.org; and GraphPad Prism were used for statistical analyses. Summary statistics are provided for demographic characteristics shown in Table 1. Continuous variables shown as mean ± standard deviation (SD). Categorical variables are shown as counts. Univariate linear regressions were performed with MG-adducts as the response variable and each of the covariates was put in the model one at a time, for AA/B and EA respectively. The set of the covariates includes age, BMI, prostate cancer (yes/no), education (high school or below/above high school), smoking (yes/yes, but quit/no), alcohol (yes/yes,but quit/no), married (yes or lived as married/no), family cancer history (yes/no). Backward stepwise regression was used to build the multivariate model.

**Table 1.**
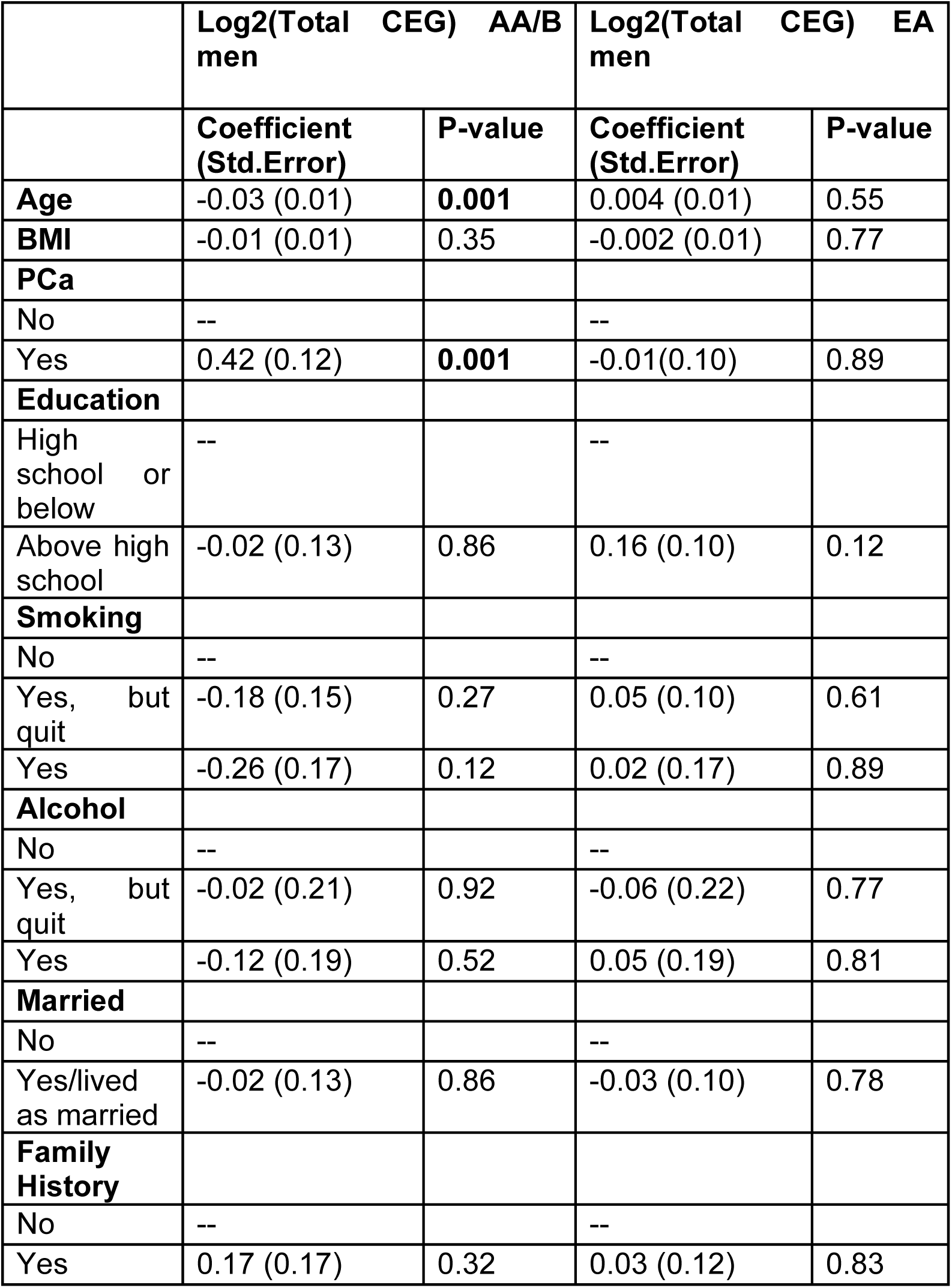
CEG is significantly associated with age and PCa status in AA/B men. Univariate linear regression analysis was performed.

|  | Log2(Total CEG) AA/B men |  | Log2(Total CEG) EA men |  |
| --- | --- | --- | --- | --- |
|  | Coefficient (Std.Error) | P-value | Coefficient (Std.Error) | P-value |
| **Age** | -0.03 (0.01) | **0.001** | 0.004 (0.01) | 0.55 |
| **BMI** | -0.01 (0.01) | 0.35 | -0.002 (0.01) | 0.77 |
| **PCa** |  |  |  |  |
| No | -- |  | -- |  |
| Yes | 0.42 (0.12) | **0.001** | -0.01(0.10) | 0.89 |
| **Education** |  |  |  |  |
| High school or below | -- |  | -- |  |
| Above high school | -0.02 (0.13) | 0.86 | 0.16 (0.10) | 0.12 |
| **Smoking** |  |  |  |  |
| No | -- |  | -- |  |
| Yes, but quit | -0.18 (0.15) | 0.27 | 0.05 (0.10) | 0.61 |
| Yes | -0.26 (0.17) | 0.12 | 0.02 (0.17) | 0.89 |
| **Alcohol** |  |  |  |  |
| No | -- |  | -- |  |
| Yes, but quit | -0.02 (0.21) | 0.92 | -0.06 (0.22) | 0.77 |
| Yes | -0.12 (0.19) | 0.52 | 0.05 (0.19) | 0.81 |
| **Married** |  |  |  |  |
| No | -- |  | -- |  |
| Yes/lived as married | -0.02 (0.13) | 0.86 | -0.03 (0.10) | 0.78 |
| **Family History** |  |  |  |  |
| No | -- |  | -- |  |
| Yes | 0.17 (0.17) | 0.32 | 0.03 (0.12) | 0.83 |

Additional statistical analyses were performed using GraphPad Prism 10.5.0 (GraphPad Software; RRID:SCR_002798. Data are shown as means ± standard deviation (SD) as indicated in the figure legends. Statistical analyses of three or more groups were compared using one- or two-way ANOVA with Tukey, Dunnett, or Brown-Forsythe multiple comparisons tests. Unpaired t-tests were used when performing statistical analyses of two groups. Statistical analyses are outlined within the figure legend. P values are notated with asterisk * p < 0.05, ** p < 0.01, *** p < 0.001, and **** p < 0.0001; 95% Confidence interval of difference. Each experiment was performed in technical duplicates or triplicates. Biological replicates for each experiment are indicated in the figure legends. For transcriptomic analysis, differential gene expression p-values were calculated using the Wald-test, followed by multiple test corrections using the Benjamini-Hochberg method. In pathway analysis, p-values were calculated using a right-tailed Fisher’s Exact Test and similarly corrected for multiple testing using the Benjamini-Hochberg method. Pathway activation or inhibition was predicted using z-scores and were calculated by the IPA software as previously described (PMID: 24336805).

### Data Availability

The data generated in this study are available from the corresponding author upon request. The RNA-sequencing data have been deposited at the NCBI Gene Expression Omnibus (GEO) under accession number GSE310499.

## Results

### MG-adducts and sRAGE are significantly altered in AA/B men with PCa

To determine the relationship of MG-adducts to PCa incidence and race, we accessed serum samples collected by Dr. Rick Kittles from AA/B and EA men from the Chicago area. We designed a nested case-control study comprised of AA/B men with (n = 83) and without (n = 83) PCa and EA men with (n = 77) and without (n = 128) PCa (Figure 1A). Serum samples were spiked with isotopically labeled standards of CEG and CEdG. Labeled and unlabeled analytes enriched using solid phase extraction were analyzed using stable isotope dilution LC-MS/MS. CEdG was not detectable in the serum samples, but CEG was observed. AA/B men without PCa had an average CEG of 1.85 ± 0.80 pmol/mL, AA/B with PCa 2.46 ± 1.33, EA without PCa 2.00 ± 0.89 and EA with PCa 1.98 ± 0.78 (Figure 1B). We found that CEG was significantly elevated in AA/B men with PCa compared to AA/B or white men of either group (Figure 1B). Univariate analysis revealed that two variables were associated with CEG in AA/B men: age and PCa status (Table 1). Multivariate logistic regression found that CEG remained significantly associated with PCa in AA/B men when controlling for age or alcohol consumption (Table 2). In EA men, no variable showed a significant association with CEG (Table 1). To further understand the potential biological implications of the elevated MG-adducts, we measured serum sRAGE, which acts as a scavenger for extracellular MG-adducts. AA/B men without PCa had an average sRAGE of 722 ± 439.5 pg/mL; AA/B with PCa, 659.7 ± 440.8 pg/mL; EA without PCa 917.5 ± 415.2 pg/mL; and EA with PCa 995.8 ± 546.8 pg/mL (Figure 1C). We discovered that sRAGE was significantly lower in AA/B men who develop PCa compared to EA or AA/B men in either group (Figure 1C). We next calculated the ratio of CEG to sRAGE. AA/B men without PCa had an average ratio of 143.7 ± 118.5; AA/B with PCa 173.1 ± 123.1; EA without PCa 79.65 ± 41.48; and EA with PCa 81.22 ± 40.32. AA/B men with and without PCa both had a significantly higher CEG/sRAGE ratio when compared to EA with or without PCa (Figure 1D). A higher ratio may indicate more circulating MG-adducts that are free to bind and activate the RAGE receptor.

**Figure 1.**
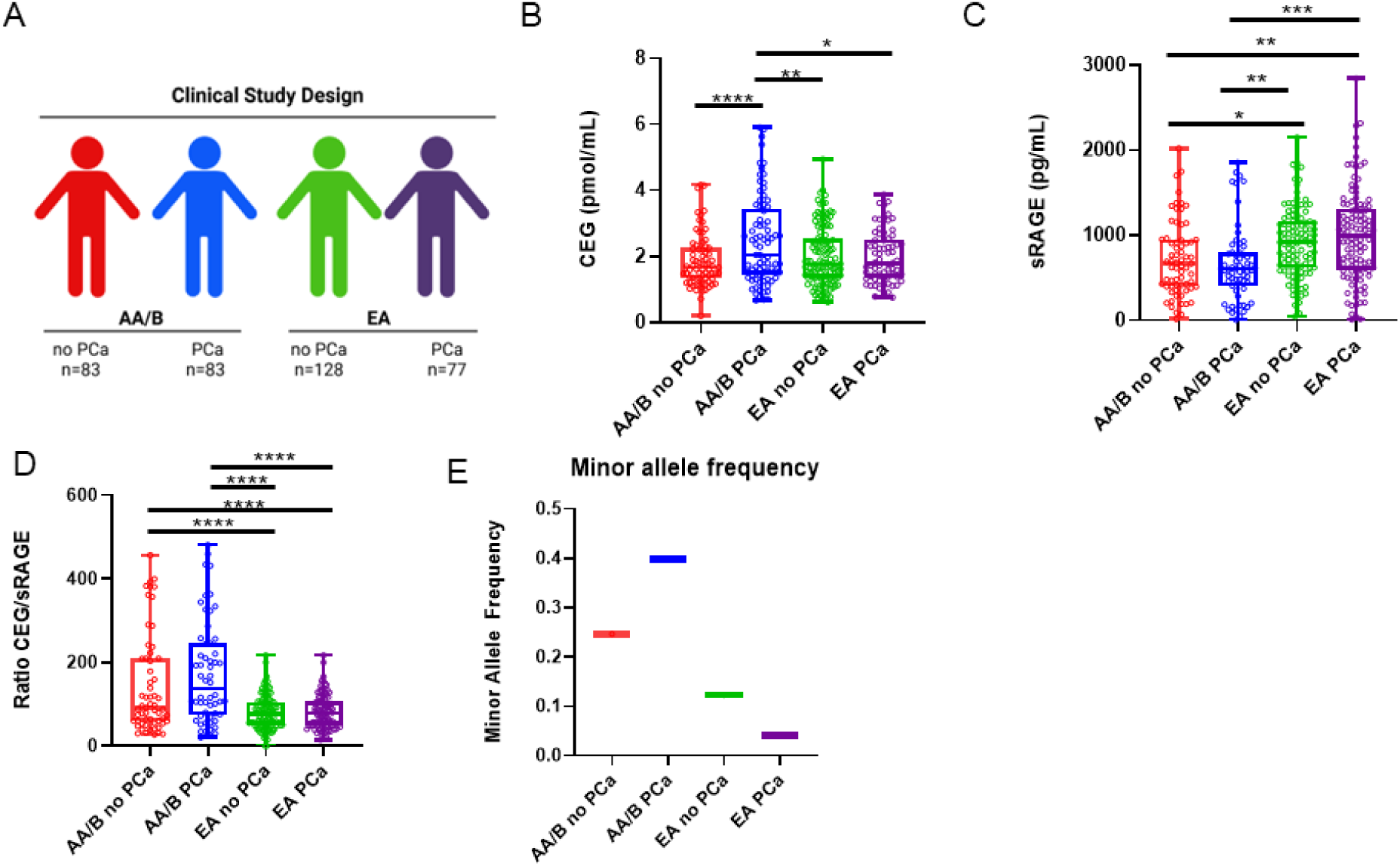
MG-adducts and sRAGE are differentially expressed in AA/B men with PCa. **A,** Nested case-control study designed from a larger clinical cohort of AA/B men with (n=83) or without (n=83) PCa and EA men with (n=77) and without (n=128) PCa. **B,** Serum levels of CEG measured using LC-MS/MS. **C,** Serum sRAGE measured using ELISA. **D,** Ratio of serum CEG to serum sRAGE. **E,** Minor allele frequency of rs1049346 in circulating genomic DNA. Significance was determined using one-way ANOVA. * p < 0.05, ** p < 0.01, *** p < 0.001, **** p < 0.0001.

**Table 2.**
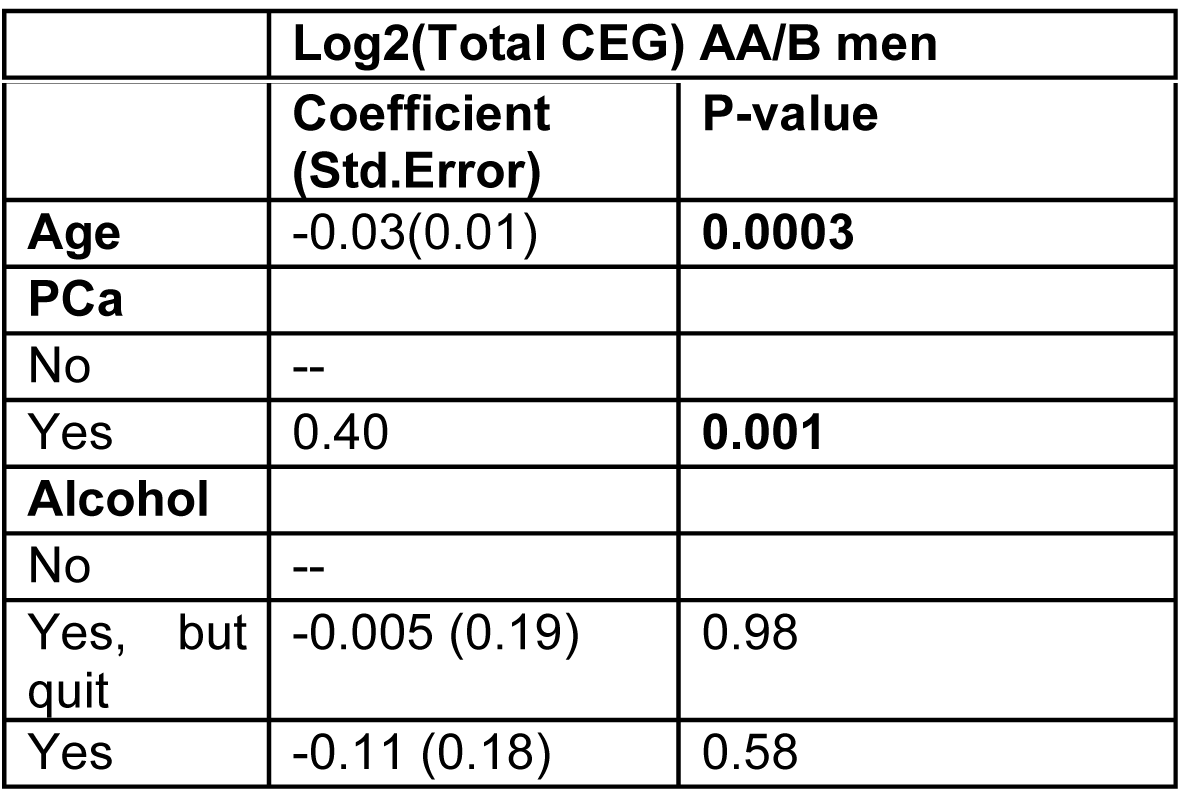
CEG remains significantly associated with PCa in AA/B men when other variables are included. Multivariable logistic regression of CEG and clinical variables with PCa.

|  | **Coefficient**<br>**(Std.Error)** | **P-value** |
| --- | --- | --- |
| **Age** | -0.03(0.01) | **0.0003** |
| **PCa** |  |  |
| No | -- |  |
| Yes | 0.40 | **0.001** |
| **Alcohol** |  |  |
| No | -- |  |
| Yes, but<br>quit | -0.005 (0.19) | 0.98 |
| Yes | -0.11 (0.18) | 0.58 |

### Single nucleotide polymorphisms in the GLO1 gene are associated with PCa status in AA/B men

To understand the role of the primary MG-detoxifying enzyme GLO1 in PCa health disparities, we determined the expression of previously characterized GLO1 SNPs in patients from our clinical study. We also performed an untargeted SNP analysis to identify novel polymorphisms that may be present. We identified multiple SNPs that were associated with self-identified race or cancer, but only one that showed significance for both self-identified race and cancer, rs1049346 (Figure 1).

### PCa cells from an AA/B man are homozygous for rs1049346 and have decreased GLO1 mRNA and protein expression compared to PCa cells from an EA man

We next characterized GLO1 expression in PCa cell lines derived from AA/B or EA men. MDA-PCa-2b are PCa cells derived from an AA/B male while C4-2 are PCa cells are derived from an EA male. Analysis of GLO1 SNP rs1049346 in these cell lines revealed that the C>T variant was heterozygous in C4-2 cells but homozygous in MDA-PCa-2b cells (Figure 2A). Moreover, we found that MDA-PCa-2b cells expressing the rs1049346 SNP had lower GLO1 mRNA and protein expression and activity compared to C4-2 cells (Figure 2B-E). To mimic hyperglycemic conditions present in diabetes, C4-2 and MDA-PCa-2b cells were grown in 7 mM glucose (normoglycemic conditions) and then treated with 25 mM glucose (hyperglycemic conditions). To induce stress associated with hyperglycemia, cells were treated with 500 µM MG. C4-2 cells treated with 25 mM glucose or 500 µM MG showed a slight increase in GLO1 protein expression while MDA-PCa-2b did not show any change (Figure 2B-C). Intriguingly, MDA-PCa-2b had significantly lower GLO1 protein expression compared to C4-2 cells in all tested conditions (Figure 2E).

**Figure 2.**
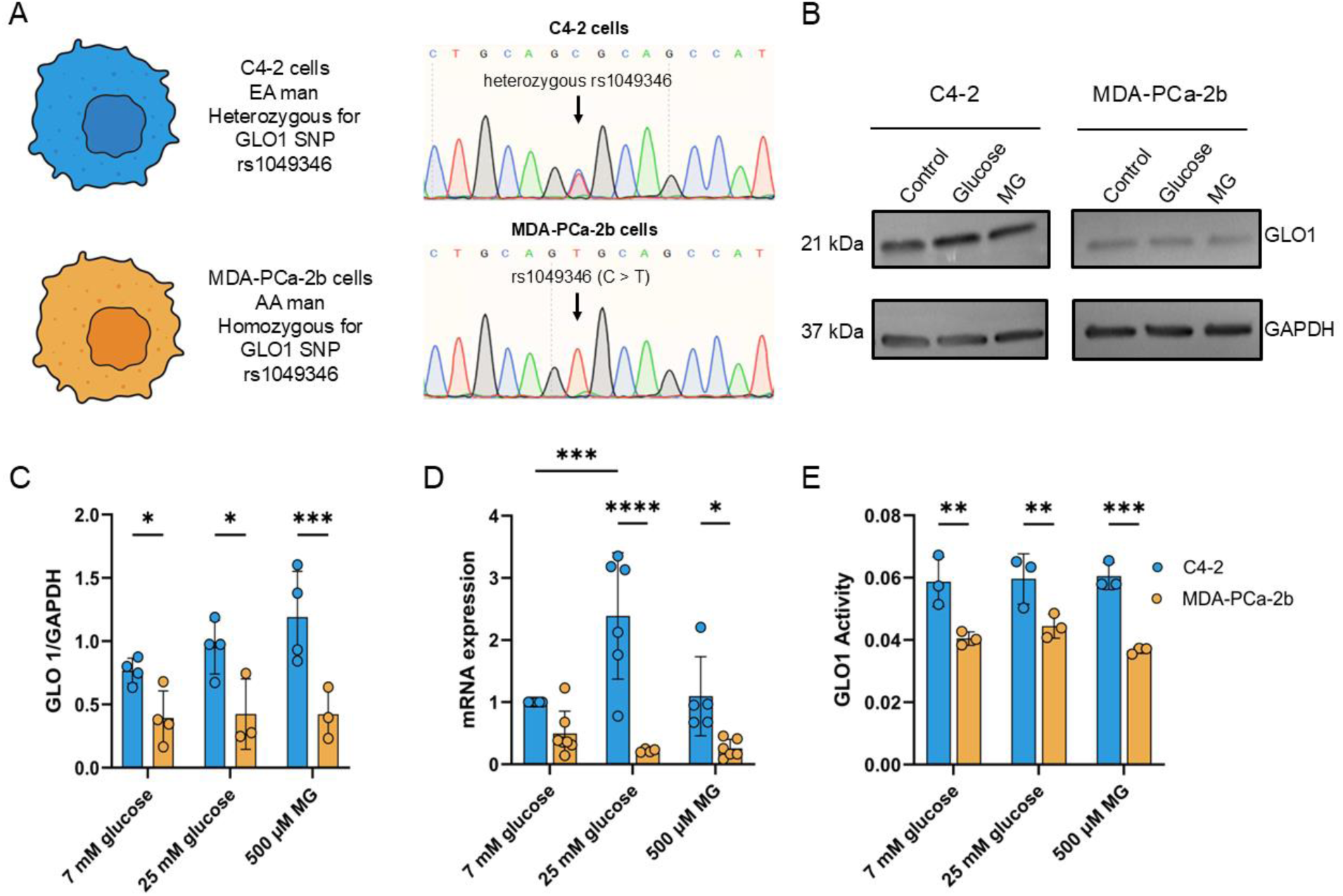
PCa cells derived from an AA/B man have decreased GLO1 mRNA and protein expression, and decreased GLO1 activity. **A,** GLO1 SNP rs1049346 was sequenced using Sanger sequencing. **B,** Western blot images of C4-2 and MDA-PCa-2b protein lysates probed for GLO1 protein expression with GAPDH as an internal control. **C,** Western blot quantification of data in B. A total of n = 4 biological replicates were completed for analysis for C4-2 basal and experimental conditions and basal MDA-PCa-2b and n = 3 biological replicates were completed for the hyperglycemic and high MG treatments for MDA-PCa-2b. **D,** qPCR analysis of GLO1 mRNA under each specified condition with experimental conditions normalized to basal C4-2 cell line conditions (7 mM glucose). A total of n = 6 biological replicates were completed in technical triplicate and analyzed using two-way ANOVA. E. GLO1 Activity was measured in protein lysates. A total of n = 3 biological replicates were completed in technical triplicate and analyzed using an unpaired t-test.

### MDA-PCa-2b cells have increased accumulation of MG-adducts compared to C4-2 cells

To determine the cellular impact of reduced GLO1 activity, we measured the accumulation of MG-adducts in DNA and RNA using LC-MS/MS. We discovered that in C4-2 cells, which have higher GLO1 activity than MDA-PCa-2b cells, stimulation with glucose or MG did not change CEdG or CEG levels (Figure 3A, 3C). However, in MDA-PCa-2b cells, which have lower GLO1 activity, treatment with glucose or MG significantly increased the levels of CEdG and CEG (Figure 3A, 3C). To determine if confounding factors may be associated with MG-adduct formation, we analyzed differentially expressed genes involved in RNA stability or DNA repair. Expression of RNA processing genes were altered by 500 µM MG treatment, with MDA-PCa-2b cells exhibiting 14 upregulated and 14 downregulated genes, whereas C4-2 cells showed a broader response, with 29 upregulated and 28 downregulated genes (Figure 3B). In MDA-PCa-2b cells, these genes include downregulation of splicing and transcription initiation factors and upregulation of mRNA surveillance and decay components. For DNA repair genes, MDA-PCa-2b cells treated with 500 µM MG exhibited greater gene expression (12 upregulated genes and 21 down regulated genes) compared to C4-2 cells treated with 500 µM MG (3 upregulated genes, 10 downregulated genes) (Figure 3D). Genes downregulated in MDA-PCa-2b include those involved with double strand break repair and base excision repair. Glucose does not significantly change gene expression in either DNA gene repair or RNA processing, which may reflect changes in acute (MG) versus chronic (glucose) exposure.

**Figure 3.**
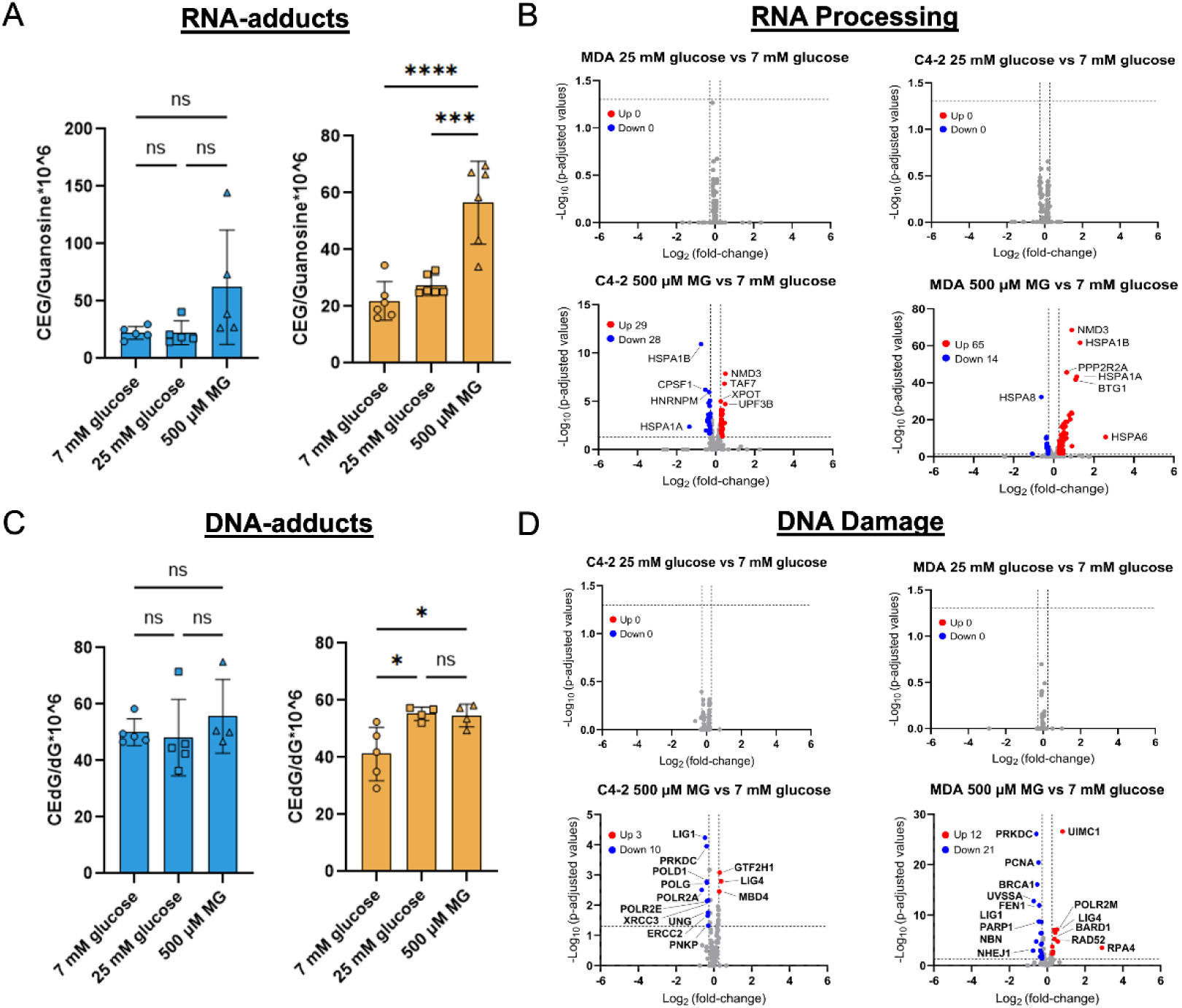
MDA-PCa-2b cells have increased accumulation of MG-adducts and altered expression of genes involved in DNA repair and RNA processing compared to C4-2 cells. CEG **A,** and CEdG **C,** were quantified using stable isotope dilution LC-MS/MS. Cells were treated with normoglycemia (7 mM glucose), hyperglycemia (25 mM glucose), or 500 μM MG. A total of n = 6 biological replicates were completed for CEG analysis in MDA-PCa-2b cells and n = 5 biological replicates were completed for CEdG analysis in MDA-PCa-2b cells and CEG and CEdG analysis in C4-2. **B,** Volcano plot presentation of genes identified to be involved in RNA processing from KEGG pathway analysis. **D,** Volcano plot of gene expression changes in genes involved in DNA damage response. * p < 0.05, ** p< 0.01, *** p < 0.001, **** p < 0.0001. Data represent n = 3 biological replicates for volcano plot analysis.

### Metabolic stress induces changes in DNA repair protein expression

In addition to the accumulation of MG-adducts induced by reduced GLO1 activity, adducts may also be persistent due to decreased DNA repair capacity. To determine the impact of metabolic stress on DNA repair protein expression, we performed a Luminex DNA Damage 7-Plex assay on protein lysates isolated from C4-2 and MDA-PCa-2b cells grown in normoglycemic (7 mM glucose), hyperglycemic (25 mM glucose), or MG (500 μM) conditions. Analysis of phosphorylated Chk1, Chk2, H2A.X, and p53 and total MDM2, p21, and ATR proteins were evaluated. Several DNA damage markers were increased, including phosphorylation of Chk1, Chk2, and H2A.X in MDA-PCa-2b cells treated with MG, but not in C4-2 cells (Figure 4A). Furthermore, proteins associated with modulating DNA repair such as ATR, p53, MDM2, and p21 were downregulated in MDA-PCa-2b compared to C4-2 (Figure 4A). We then analyzed differences between cell lines and treatments for each individual protein. MDA-PCa-2b cells had significantly lower total ATR protein expression compared to C4-2 cells when both were grown in hyperglycemic conditions (Figure 4B). Analysis of DNA damage markers revealed significantly higher Chk1 phosphorylation in MDA-PCa-2b in all conditions compared to C4-2 cells. Additionally, MG treatment significantly increased Chk1 phosphorylation compared to 7 and 25 mM glucose treatment (Figure 4C). MDA-PCa-2b cells treated with MG had significantly increased Chk2 phosphorylation compared to 7 or 25 mM glucose (Figure 4C). Comparing C-42 to MDA-PCa-2b revealed a significant difference between basal and glucose conditions but were not different between cell lines (Figure 4C). MDA-PCa-2b cells treated with MG had a significant increase in phosphorylated H2A.X(Figure 4D). Additionally, MDA-PCa-2b cells had significantly lower p21 expression, a marker of DNA repair activation, compared to C4-2 cells in all conditions tested (Figure 4D).

**Figure 4.**
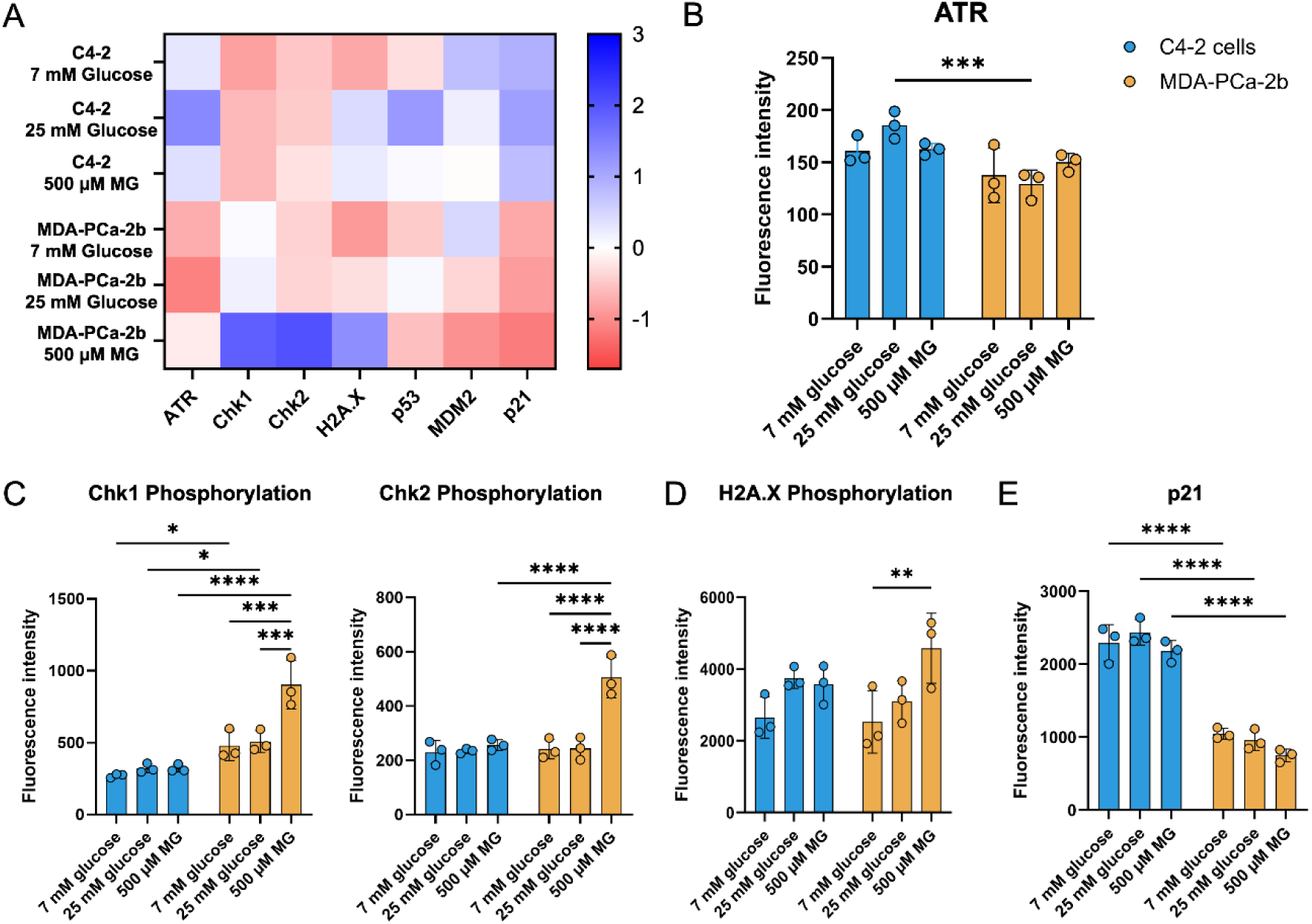
Metabolic stress induces differential expression of DNA repair proteins in MDA-PCa-2b cells. **A,** Heatmap of the Luminex assay data shows trends in various proteins associated with DNA damage and repair. **B,** Analysis of ATR protein expression. **C,** Phosphorylated Chk1 and Chk2 protein expression. **D,** Phosphorylated H2AX protein expression. **E,** p21 protein expression. A total of n=3 biological replicates in technical duplicate were completed for both MDA-PCa-2b and C4-2 cells. Two-way ANOVA with Tukey’s multiple comparisons test was utilized to compare analogous conditions within a cell line. * p < 0.05, ** p< 0.01, *** p < 0.001, **** p < 0.0001.

### Acute MG exposure reprograms stress response gene expression in PCa cells

We next investigated the impact of MG stress on gene expression in C4-2 and MDA-PCa-2b cells. Cells were treated with 7 mM or 25 mM glucose for 24 hours, or 500 µM MG for 3 hours and then RNA-Seq was performed. Principal Component Analysis (PCA) revealed distinct clustering by cell line (PC1, 39.76%) and separation of MG-treated samples from glucose conditions (PC2, 10.27%) (Figure 5A). Minimal separation between 25 mM and 7 mM glucose suggested that hyperglycemia induced minimal transcriptomic changes. Differential expression analysis (FDR < 0.05, |fold-change| ³ 2) revealed no significant differentially expressed genes (DEGs) in response to 25 mM glucose in MDA-PCa-2b cells (Figure 5B), and only 2 DEGs in C4-2 cells (*GAGE12* and *GAGE13*) (Figure 5C). In contrast, 500 µM MG induced transcriptomic changes in 384 upregulated and 136 downregulated DEGs in MDA-PCa-2b, and 96 upregulated and 8 downregulated in C4-2 (Figure 5B and C, respectively). We next characterized genes uniquely differentially regulated with MG treatment in MDA-PCa-2b and C4-2 cells. We found that MG induced 463 unique DEGs in MDA-PCa-2b, 46 unique in C4-2, and 58 shared between both cells (Figure 5D). Upregulated DEGs unique to MDA-PCa-2b include *TXNIP, RGS2, NR3C1, JUN, HMOX1*, *CXCR4*, and *MT1G.* Intriguingly, TXNIP is inversely related to GLO1 expression (PMID: 33360689) while the others are associated with redox regulation, stress response, metastasis, and ferroptosis resistance. Upregulated C4-2 specific genes include *ATF4, CLIC4, IGF2BP2*, and *EGR2* and are linked to ER stress, apoptosis, and differentiation. Both cell lines shared common upregulated DEGs (*CHAC1, DDIT3, ATF3, GDF15, SESN2*) and reflect core MG stress responses involving the unfolded protein response and oxidative stress adaptation (PMID: 35635155).

**Figure 5.**
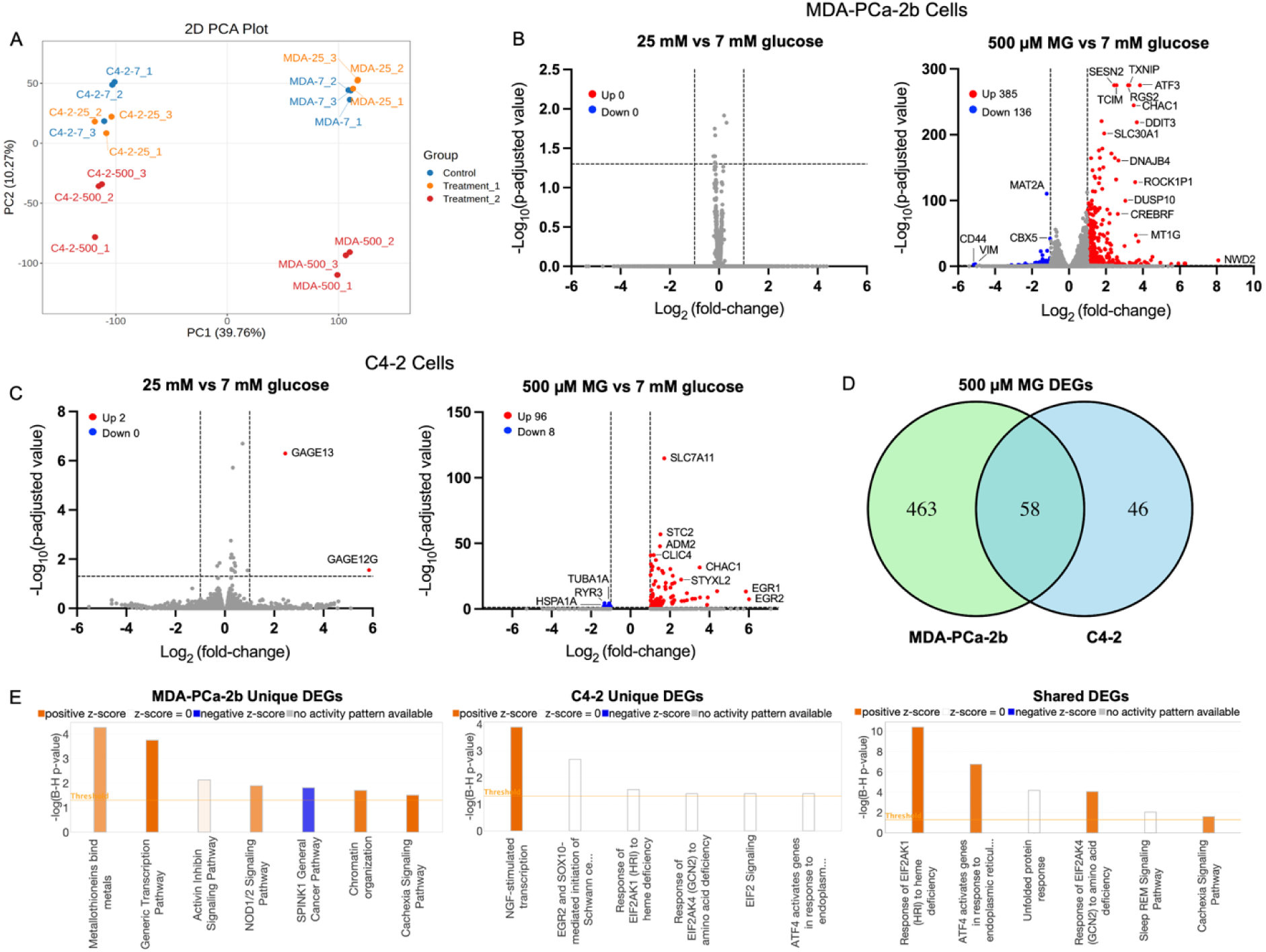
Transcriptomic analysis of C4-2 and MDA-PCa-2b cells in response to glucose and MG treatments. **A,** Principal component analysis showing global transcriptomic variation in 7 mM glucose (blue), 25 mM glucose (orange) and 500 µM MG (red) in both cell lines. **B,** Volcano plots showing DEGs comparing 25 mM vs 7 mM glucose (left) and 500 µM MG vs 7 mM glucose (right) in MDA-PCa-2b cells. **C,** Same comparison in C4-2 cells. Significance was determined by –Log_10_(p-adjusted value) > 1.3 and |Log_2_ (fold-change)| ≥ 1, where blue indicates downregulated and red indicates upregulated. **D,** Venn diagram showing overlap of DEGs in response to 500 µM MG vs 7 mM glucose between MDA-PCa-2b (green) and C4-2 (blue). **E,** Ingenuity Pathway Analysis of significantly enriched canonical pathways among MG-responsive genes unique to MDA-PCa-2b (left), C4-2 (center), or shared between both cells (right). Orange bars indicate predicted activation (z-score > 0), blue indicates predicted inactivation (z-score < 0) and white indicates no predicted activity (z-score = 0). Data represent n = 3 biological replicates.

Ingenuity Pathway Analysis (IPA) was performed on shared and uniquely identified DEGs. This analysis revealed seven significantly enriched pathways in MDA-PCa-2b cells, including Metallothionein, Activin/Inhibin, Chromatin organization, and Cachexia signaling (Figure 5E). In C4-2 cells, while several pathways were significantly enriched, only NGF-stimulated transcription was activated (Figure 5E). Six pathways were enriched in both cell lines, such as the Unfolded Protein Response and ATF4-mediated ER stress, indicating a conserved transcriptomic response to MG (Figure 5E).

### Hyperglycemia and MG increase rates of oxidative phosphorylation and glycolysis with associated gene expression changes in MDA-PCa-2b but not C4-2 cells

We utilized the ATP Rate Seahorse Assay (Agilent) to determine the impact of environmental stressors within each cell line. In hyperglycemic conditions, MDA-PCa-2b cells had a pronounced increase rate of oxidative phosphorylation while high MG conditions significantly decreased mitochondrial metabolism (Figure 6A). Hyperglycemia increased glycolysis in MDA-PCa-2b (Figure 6C). C4-2 cells did not show a significant change in either metabolic pathway when subjected to hyperglycemia or MG (Figure 6A, C). Transcriptomic analysis indicates that there is upregulation of a greater number of genes associated with processes related to glycolysis and oxidative phosphorylation in MDA-PCa-2b cells treated with MG but minimally affected in C4-2 cells supporting the possibility that MG affects gene expression in MDA-PCa-2b differentially compared to C4-2 cells.

**Figure 6.**
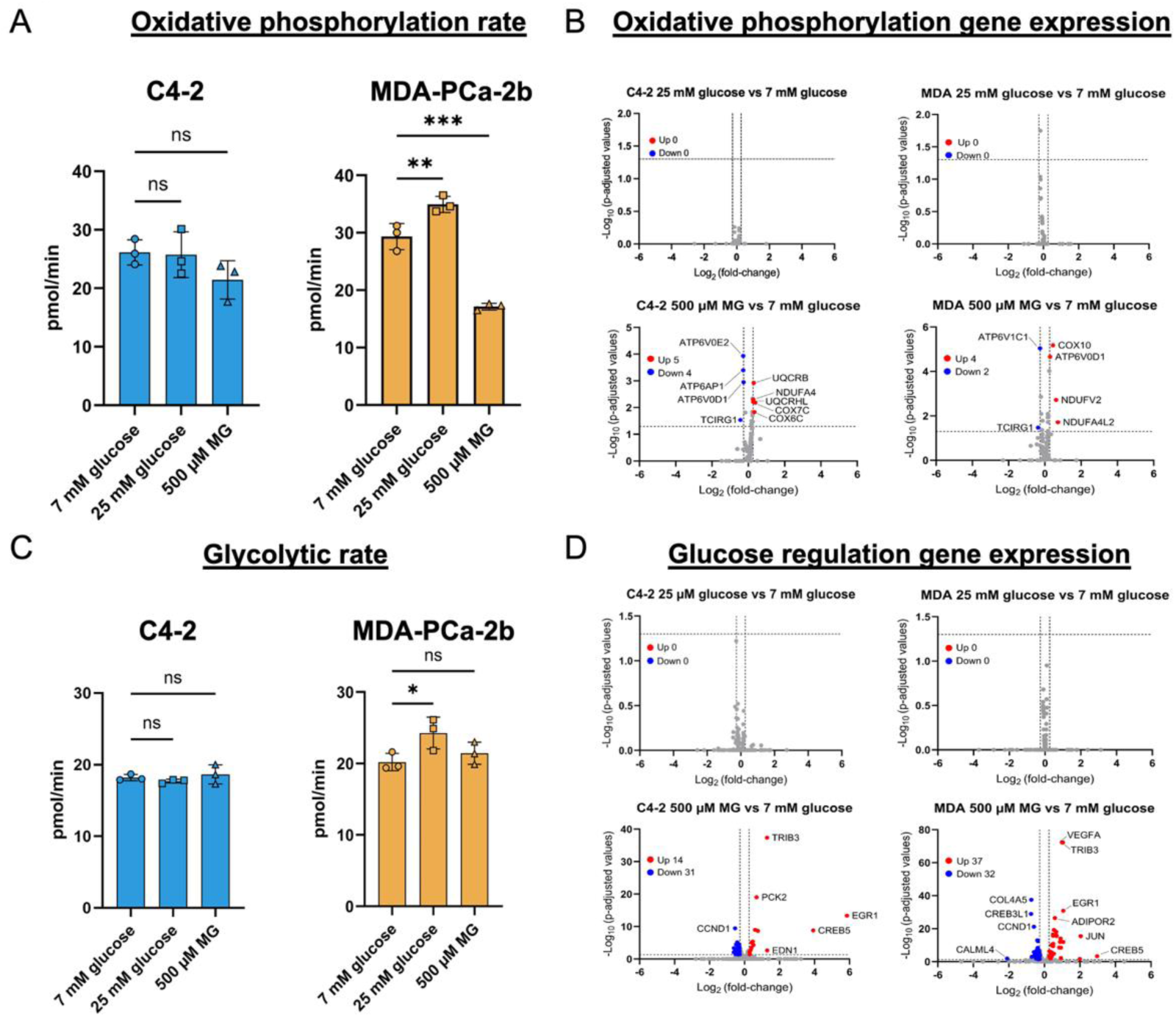
Hyperglycemia and MG alter oxidative phosphorylation and glycolytic rates in MDA-PCa-2b cells. Agilent Seahorse XeF96 technology was used to determine the rate of oxidative phosphorylation **A,** and glycolysis **C,** when C4-2 and MDA-PCa-2b cells grown in 7 mM glucose were treated with 25 mM glucose or 500 μM MG. One-way ANOVA was used to compare treatments within a cell line. Expression of genes involved in oxidative phosphorylation **B,** or glucose regulation **D,** were analyzed using volcano plots with comparisons between 7 mM glucose and 25 mM glucose or 7 mM glucose and 500 μM MG, as indicated. * p < 0.05, ** p < 0.01, *** p < 0.001. Oxidative phosphorylation and glycolysis were analyzed in technical triplicate and n = 3 biological replicates for each condition measured. For volcano plots, data represent n = 3 biological replicates.

### MDA-PCa-2b cells have increased rates of proliferation when subjected to hyperglycemic and high MG conditions

As changes in metabolic processes and DNA adduct accumulation may impact cell proliferation, we determined the cell number in each cell line treated with glucose or MG. We discovered that MDA-PCa-2b cells grown in normoglycemic and hyperglycemic conditions had increased proliferative capacity than C4-2 cells (Supplementary Figure 1).

## Discussion

PCa outcomes differ across patient groups, including AA/B men, individuals with obesity, and those with diabetes.^17–19^ In this study, we found that RNA MG-adducts are elevated in AA/B men with PCa and are associated with increased frequency of the rs1049346 GLO1 5’-UTR variant. Using PCa cells from an AA/B man and cells from an EA man, we characterized how different racial background coupled with expression of rs1049346 impacted the response of cells to metabolic stress. In PCa cells derived from an AA/B man, GLO1 activity was lower and MG-induced DNA and RNA adducts were higher than in cells from an EA man, and the two models showed distinct transcriptional responses to metabolic stress.

A central insight is the regulatory potential of rs1049346, which lies within a putative NRF2-recognition site, an antioxidant response element (ARE) (sequence GTGATAGCA), located within the GLO1 5’-UTR.^20^ Indeed, as a master regulator of oxidative stress response, GLO1 is a major target gene of NRF2 and NRF2 activation reduces MG.^20, 21^ Moreover, NRF2 controls the expression of genes involved in glutathione metabolism, a cofactor essential for GLO1 activity.^22^ Therefore, as rs1049346 may weaken NRF2 binding to the GLO1 promoter region, this could reduce GLO1 transcription and activity. This mechanism aligns with the lower GLO1 expression and higher MG-adduct burden observed in cells from an AA/B man. TXNIP is a well-established MG-responsive gene that increases when NRF2-driven antioxidant capacity is low.^23^ Indeed, NRF2 directly binds an ARE located upstream to TXNIP promoter to downregulate its expression under basal and high glucose.^24^ Thus, TXNIP upregulation in MG-treated in MDA-PCa-2b cells compared to C4-2 cells may reflect an overall inhibition of NRF2 activity in these cells. Future studies should further explore the role of NRF2 signaling in obesity-associated PCa progression.

MG accumulation affected multiple processes relevant to PCa biology, including DNA repair, RNA processing, oxidative stress, and cellular metabolism. MG-RNA adducts destabilize transcripts and may account for increased RNA-surveillance and processing pathways.^4^ MG exposure also enhanced glycolysis, reduced oxidative phosphorylation, and activated metallothionein signaling and NR3C1 in MDA-PCa-2b cells, whereas C4-2 cells mounted a more limited response. Hyperglycemia alone produced minimal transcriptional changes, suggesting that glucose-induced differences between these lines arise primarily from post-translational regulation rather than transcriptional reprogramming. The overall impact of these changes resulted in increased cell proliferation in MDA-PCa-2b cells compared to C4-2 cells.

While our work provides a compelling description of the relationship between glycolytic stress, race, and altered detoxification capacity, a limitation of our work is the use of non-isogenic cell lines, which prevents definitive attribution of phenotypes to rs1049346. In this work, our goal was to determine differences between PCa cells from different racial backgrounds, but future work will incorporate isogenic cell lines to determine the direct impact of GLO1 SNPs on MG-adduct accumulation in these models.

In summary, our results support a model in which the rs1049346 5’-UTR variant reduces GLO1, decreases glyoxalase activity, and increases MG-adduct accumulation and downstream stress responses. These findings highlight glycation stress as a potential contributor to PCa progression in metabolic dysfunction and support therapeutic strategies that target diabetes-related metabolic stress in individuals with PCa.

## Authors’ Contributions

**A. Battad:** Formal analysis, investigation, visualization, methodology, writing original draft, and editing. **F.J. Nunez:** Formal analysis, investigation, visualization, methodology, writing-original draft, and editing. **M. Walker:** Formal analysis, investigation, and editing. **K. Sun:** Formal analysis, investigation, and editing. **E. Desjarlais:** Formal analysis, investigation, and editing. **S. Hodge:** Formal analysis, investigation, and editing. **E.D.J.L. Gonzales:** Formal analysis, investigation, and editing. **S.W.T. Lai**: Formal analysis, investigation, and editing. **L. Woods-Burnham:** Formal analysis, investigation, visualization, methodology, and editing. **S. Hooker:** Formal analysis, investigation, visualization, methodology, and editing. **R. Aburajab:** Formal analysis, investigation, visualization, methodology, and editing. **Q. Feng:** Formal analysis, investigation, visualization, methodology, and editing. **M. Talley:** Formal analysis, investigation, visualization, methodology, and editing. **S. Baumel-Alterzon**: Formal analysis, visualization, and editing. **D.W. Craig:** Formal analysis and editing. **R.A. Kittles:** Formal analysis, investigation, visualization, methodology, and editing. **Y.R. Li:** Formal analysis, investigation, visualization, methodology, and editing. **J. Termini:** Formal analysis, investigation, visualization, methodology, and editing. **S.C. Shuck:** Formal analysis, investigation, visualization, methodology, writing original draft, and editing.

## Ethics statement

This study was conducted in accordance with the Declaration of Helsinki and approved by the Institutional Review Board. Written informed consent was obtained from all participants, and all human biospecimens were collected and used following institutional guidelines and regulatory requirements. All human biospecimens were de-identified prior to receipt, and the study was determined by the IRB to be exempt from human subjects research regulations.

## Acknowledgements

S.C.S. received research support from NIH R21CA282612 with R21CA282612-01S1 to K.S.. Author L.W.-B. received research support from Georgia CTSA NIH KL2TR002381 and NIH UL1 TR002378, Department of Defense Prostate Cancer Research Program W81XWH2110038, and Prostate Cancer Foundation 20YOUN04. Authors L.W.-B. and R.A.K. received research support from NIH/NIMHD 2U54MD007602. Work was performed in collaboration with the Integrated Mass Spectrometry and Integrated Genomics core facilities at City of Hope, which are supported by NCI grant P30CA03357.

## References

1. Saka, A. H.; Giaquinto, A. N.; McCullough, L. E.; Tossas, K. Y.; Star, J.; Jemal, A.; Siegel, R. L., Cancer statistics for African American and Black people, 2025. 2025, 75 (2), 111–140.

2. Bancks, M. P.; Kershaw, K.; Carson, A. P.; Gordon-Larsen, P.; Schreiner, P. J.; Carnethon, M. R., Association of Modifiable Risk Factors in Young Adulthood With Racial Disparity in Incident Type 2 Diabetes During Middle Adulthood. Jama 2017, 318 (24), 2457–2465.

3. Lai, S. W. T.; Lopez Gonzalez, E. J.; Zoukari, T.; Ki, P.; Shuck, S. C., Methylglyoxal and Its Adducts: Induction, Repair, and Association with Disease. Chemical research in toxicology 2022, 35 (10), 1720–1746.

4. Lopez Gonzalez, E. J.; Tsuen Lai, S. W.; Sun, K.; Carson, C. R.; Hernandez-Castillo, C.; Zoukari, T.; Lopez, K.; Zhang, J.; Blevins, T.; Termini, J.; Shuck, S. C., Methylglyoxal-induced RNA modifications decrease RNA stability and translation and are associated with type 2 diabetes. Molecular metabolism 2025, 102186.

5. Shuck, S. C.; Wuenschell, G. E.; Termini, J. S., Product Studies and Mechanistic Analysis of the Reaction of Methylglyoxal with Deoxyguanosine. Chemical research in toxicology 2018, 31 (2), 105–115.

6. Thornalley, P. J., Pharmacology of methylglyoxal: formation, modification of proteins and nucleic acids, and enzymatic detoxification--a role in pathogenesis and antiproliferative chemotherapy. General pharmacology 1996, 27 (4), 565–73.

7. Romanuik, T. L.; Ueda, T.; Le, N.; Haile, S.; Yong, T. M.; Thomson, T.; Vessella, R. L.; Sadar, M. D., Novel biomarkers for prostate cancer including noncoding transcripts. The American journal of pathology 2009, 175 (6), 2264–76.

8. Rounds, L.; Nagle, R. B.; Muranyi, A.; Jandova, J.; Gill, S.; Vela, E.; Wondrak, G. T., Glyoxalase 1 Expression as a Novel Diagnostic Marker of High-Grade Prostatic Intraepithelial Neoplasia in Prostate Cancer. Cancers 2021, 13 (14).

9. Zeng, Q.; Yang, T.; Wei, W.; Zou, D.; Wei, Y.; Han, F.; He, J.; Huang, J.; Guo, R., Association between GLO1 variants and gestational diabetes mellitus susceptibility in a Chinese population: a preliminary study. Frontiers in endocrinology 2023, 14, 1235581.

10. Peculis, R.; Konrade, I.; Skapare, E.; Fridmanis, D.; Nikitina-Zake, L.; Lejnieks, A.; Pirags, V.; Dambrova, M.; Klovins, J., Identification of glyoxalase 1 polymorphisms associated with enzyme activity. Gene 2013, 515 (1), 140–3.

11. Wu, J. C.; Li, X. H.; Peng, Y. D.; Wang, J. B.; Tang, J. F.; Wang, Y. F., Association of two glyoxalase I gene polymorphisms with nephropathy and retinopathy in Type 2 diabetes. J Endocrinol Invest 2011, 34 (10), e343–8.

12. Raucci, A.; Cugusi, S.; Antonelli, A.; Barabino, S. M.; Monti, L.; Bierhaus, A.; Reiss, K.; Saftig, P.; Bianchi, M. E., A soluble form of the receptor for advanced glycation endproducts (RAGE) is produced by proteolytic cleavage of the membrane-bound form by the sheddase a disintegrin and metalloprotease 10 (ADAM10). FASEB journal : official publication of the Federation of American Societies for Experimental Biology 2008, 22 (10), 3716–27.

13. Kim, J.-Y.; Jung, J.-H.; Lee, S.-J.; Han, S.-S.; Hong, S.-H., Glyoxalase 1 as a Therapeutic Target in Cancer and Cancer Stem Cells. Molecules and Cells 2022, 45 (12), 869–876.

14. Rabbani, N., Methylglyoxal and glyoxalase 1-a metabolic stress pathway-linking hyperglycemia to the unfolded protein response and vascular complications of diabetes. Clinical science 2022, 136 (11), 819–824.

15. Bora, S.; Adole, P. S.; Vinod, K. V.; Pillai, A. A.; Ahmed, S., The genetic polymorphisms and activity of glyoxalase 1 as a risk factor for acute coronary syndrome in South Indians with type 2 diabetes mellitus. Gene 2023, 885, 147701.

16. Johnson, J. R.; Mavingire, N.; Woods-Burnham, L.; Walker, M.; Lewis, D.; Hooker, S. E.; Galloway, D.; Rivers, B.; Kittles, R. A., The complex interplay of modifiable risk factors affecting prostate cancer disparities in African American men. Nature reviews. Urology 2024, 21 (7), 422–432.

17. Leitzmann, M. F.; Ahn, J.; Albanes, D.; Hsing, A. W.; Schatzkin, A.; Chang, S. C.; Huang, W. Y.; Weiss, J. M.; Danforth, K. N.; Grubb, R. L., 3rd; Andriole, G. L., Diabetes mellitus and prostate cancer risk in the Prostate, Lung, Colorectal, and Ovarian Cancer Screening Trial. Cancer causes & control : CCC 2008, 19 (10), 1267–76.

18. Kelkar, S.; Oyekunle, T.; Eisenberg, A.; Howard, L.; Aronson, W. J.; Kane, C. J.; Amling, C. L.; Cooperberg, M. R.; Klaassen, Z.; Terris, M. K.; Freedland, S. J.; Csizmadi, I., Diabetes and Prostate Cancer Outcomes in Obese and Nonobese Men After Radical Prostatectomy. JNCI Cancer Spectrum 2021, 5 (3).

19. Feng, X.; Song, M.; Preston, M. A.; Ma, W.; Hu, Y.; Pernar, C. H.; Stopsack, K. H.; Ebot, E. M.; Fu, B. C.; Zhang, Y.; Li, N.; Dai, M.; Liu, L.; Giovannucci, E. L.; Mucci, L. A., The association of diabetes with risk of prostate cancer defined by clinical and molecular features. British journal of cancer 2020, 123 (4), 657–665.

20. Xue, M.; Rabbani, N.; Momiji, H.; Imbasi, P.; Anwar, M. M.; Kitteringham, N.; Park, B. K.; Souma, T.; Moriguchi, T.; Yamamoto, M.; Thornalley, P. J., Transcriptional control of glyoxalase 1 by Nrf2 provides a stress-responsive defence against dicarbonyl glycation. The Biochemical journal 2012, 443 (1), 213–22.

21. Nishimoto, S.; Koike, S.; Inoue, N.; Suzuki, T.; Ogasawara, Y., Activation of Nrf2 attenuates carbonyl stress induced by methylglyoxal in human neuroblastoma cells: Increase in GSH levels is a critical event for the detoxification mechanism. Biochemical and biophysical research communications 2017, 483 (2), 874–879.

22. Harvey, C. J.; Thimmulappa, R. K.; Singh, A.; Blake, D. J.; Ling, G.; Wakabayashi, N.; Fujii, J.; Myers, A.; Biswal, S., Nrf2-regulated glutathione recycling independent of biosynthesis is critical for cell survival during oxidative stress. Free radical biology & medicine 2009, 46 (4), 443–53.

23. Maimaiti, S.; Dagnell, M.; Coppo, L.; Arnér, E. S. J., Patient-derived TXNIP-deficient primary cells exhibit NRF2 activation linked to upregulation of glyoxalase 1 (GLO1). Free radical biology & medicine 2025, 239, 230–241.

24. He, X.; Ma, Q., Redox regulation by nuclear factor erythroid 2-related factor 2: gatekeeping for the basal and diabetes-induced expression of thioredoxin-interacting protein. Mol Pharmacol 2012, 82 (5), 887–97.

