## Supplemental Table 1, Table 1-9 for "Mechanistic Link Between Glyoxalase 1 Expression and Methylglyoxal-Induced Oncogenic Stress in Prostate Cancer"

Beckman Research Institute at City of Hope

Departments of

[1]Diabetes and Cancer Metabolism

[2]Population Sciences and Division of Health Equity

[3]Integrative Translational Sciences

[4]Radiation Oncology

[5]Cancer Genetics and Epigenetics

[6]Biostatistics and Mathematical Oncology

[7]Molecular and Cellular Endocrinology

[8]Cancer Biology and Molecular Medicine

*Corresponding author

Sarah C. Shuck

1500 Duarte Rd, Duarte, CA 91010

**Supplemental Methods**

*Cell Culture*

Immortalized PCa cell line C4-2 was obtained from Dr. Saul Priceman (University of Southern California) and cell line MDA-PCa-2b was obtained from Drs. Leanne Woods-Burnham and Rick Kittles (Morehouse School of Medicine). C4-2 cells were maintained in DMEM/F12K media (Innovative Research, IDMEMF120322500ML) supplemented with a final concentration of 7 mM glucose (Gibco, A2494001) and 10% FBS (Omega Scientific, FB-01). C4-2 cells between passages 10 and 21 were used for experiments. MDA-PCa-2b cells were cultured in F12K media (American Type Culture Collection, 30-2004) supplemented with 20% FBS (Omega Scientific, FB-01), and the following chemicals were from Sigma Aldrich unless otherwise indicated: 25 ng/mL cholera toxin (C8052-1MG), 10 ng/mL hEGF (E9644-.5MG), 0.002 ng/mL insulin (Fisher Scientific, 12585014), 45 nM selenious acid (211176-10g), 100 pg/mL hydrocortisone (H-0888), and 5 µM phosphoethanolamine (P0503-1G). In accordance with the National Institutes of Health (NIH) guidelines concerning the authentication of key biological resources and to ensure the identity and validity of the resource, cell lines cultured in this study were purchased from ATCC that performs cell line characterizations. Additionally, cells were authenticated utilizing Short Tandem Repeat (STR) profiling against the ATCC STR database (ATCC, Cat: ATCC 135-XV).

*Sequencing GLO1 SNPs*

C4-2 and MDA-PCa-2b cells were plated in a 6 cm dish at a density of $1 \times 10^6$ cells/plate. Approximately $5 \times 10^6$ cells were harvested and genomic DNA was extracted and purified according to manufacturer's instructions using DNeasy® Blood and Tissue Kit (Qiagen; 69506). Regions around *GLO1* SNP rs1049346 were amplified using PCR (Supplemental

Table 7). PCR products were sequenced using Sanger Sequencing (Eton Biosciences, Inc, San Diego, CA).

*Western blot*

C4-2 and MDA-PCa-2b cells were plated in 6-well plates at a density of 5 x $10^5$ cells/well. Cells were adhered overnight and treated according to the “Experimental Treatments” section outlined above. Cell pellets were harvested in 10 mM sodium phosphate buffer (J63791.AK, pH 7.0) supplemented with a Pierce protease/phosphatase inhibitor cocktail (Thermo Fisher Scientific, PIA32959) at 4 °C and sonicated at 20% amplitude for 30 s while kept on ice. Contents were then centrifuged at 17,000 x g for 30 minutes at 4 °C. Western blot analysis was performed as previously described (PMID: 38247509), with the following antibodies: GLO1 (Abcam; ab96032, 0.45 µg/mL) and GAPDH (Cell Signaling Technology, Danvers, MA, USA; 5174, 0.4 µg/mL).

*RNA Extraction and Quantitative PCR*

MDA-PCa-2b cells and C4-2 cells were plated at a density of 5x$10^5$ cells/plate in 6 cm dishes. Cells were treated according to the “Experimental Treatments” section outlined above. mRNA was isolated and cDNA was synthesized as previously described. qPCR was performed using iTaq™ Universal SYBR® Green Supermix (BioRad, 1725121) on a QuantStudio 7 Flex Real-Time PCR System (Applied Biosystems, Foster City, CA, USA). *GLO1* mRNA was quantified with forward primer 5´-CCCCAGTACCAAGGATTTTCT-3´ and reverse primer 5´- TGGGAAAATCACATTTTTGGA-3´. *GAPDH* was also quantified for normalization with forward primer 5´-TTGGCTACAGCAACAGGGTG-3´ and reverse primer 5´- GGGGAGATTCAGTGTGGTGG-3´.

*Quantitation of serum sRAGE*

sRAGE was quantified from 50 µL serum using human RAGE quantikine ELISA kit (R&D Systems) per manufacturer's instructions. Plates were read using a Synergy LX Multimode Microplate Reader (Agilent BioTek).

*GLO1 activity assay*

Please refer to the "Experimental Treatments" section above for information on treatment conditions for cells. GLO1 activity assay was performed as previously described (PMID: 24646266).

*MG Adduct Quantitation from cellular DNA and RNA*

C4-2 and MDA-PCa-2b cells grown in 7 mM glucose were plated at a density of $2x10^6$ cells/plate on 10 cm dishes and allowed to adhere overnight. Cells were treated based on the "Experimental Treatments" section. Following treatment, DNA and RNA were isolated using the DNeasy Blood and Tissue Kit (Qiagen, 69506). MG-adducts CEdG (DNA) and CEG (RNA) were quantified using LC-MS/MS as previously described (PMID:4049652).

*Luminex Assay*

C4-2 and MDA-PCa-2b cells grown in 7 mM glucose were plated at a density of $5x10^5$ cells/plate on 6 cm dishes and allowed to adhere overnight. Cells were treated as outlined in the “Experimental Treatments” section. Following treatment, protein lysates were harvested using Milliplex DNA Damage/Genotoxicity Magnetic Bead Panel supplied Cell Lysis Buffer supplemented with phosphatase/protease inhibitors (Thermo Fisher Scientific). Protein lysates were filtered with a 0.1 µm centrifuge filter (Fisher Scientific) and collected in an Eppendorf tube by centrifuging the samples at 13,000 x g for 1 minute and stored at –80 °C. Reagents and protocol for DNA Damage Assay were followed as described in manufacturer’s instructions (Millipore Sigma). Data was collected by the Analytical Pharmacology Core at City of Hope.

*Cell Counting: Trypan blue exclusion*

C4-2 and MDA-PCa-2b cells were plated in a 10 cm plate at a density of $1 \times 10^6$ cells/plate. Trypsizined cells were mixed 1:1 with trypan blue (Gibco, 8611) and live and dead cells counted using a LUNA automatic cell counter (Logos Biosystems, L40002).

*Seahorse assay: Bioenergetic profile analysis*

C4-2 and MDA-PCa-2b cells were plated at a density of 20,000 cells/well and 30,000 cells/well, respectively, in an Agilent Seahorse XFe96/XF Pro Cell Culture Microplate (103794-100). The cartridge from the manufacturer was calibrated with XF Calibrant (100840-000) and incubated in a 37 °C $CO_2$-free incubator overnight. Culture medium was removed, and cells were washed with DMEM XF base Seahorse medium (Agilent, 103575-100) twice, then placed in a 37 °C $CO_2$-free incubator for 1 hour. ATP Rate Assay was conducted on an Agilent Seahorse XFe96/XF (103591-100) by injecting 15 μM oligomycin, 7.3 μM antimycin, and 8 μM rotenone into the wells using the ATP Rate Assay protocol as provided by the manufacturer.

Table 1. Demographic characteristics of the cohort. Education was determined as 0 < high school, 1 high school, 2 some college, 3 bachelor's degree, 4 > beyond bachelor's degree. Statistical analyses – T = T-test, $\chi$ = chi-square test, LRM = logistic regression model.

*Transcriptomics*

MDA-PCa-2b and C4-2 cells were in 7 mM glucose were plated at a density of 5 x $10^5$ cells in 6 cm dishes and treated as outlined in the "Experimental Treatments" section. Total RNA was extracted using the Direct-zol RNA Miniprep Kit (Zymo Research, R2050). RNA sequencing and bioinformatic analysis were performed by Metware Biotechnology Inc. (Boston, USA) on the Illumina platform. Raw reads were quality-filtered using fastp, with removal of adapter sequences and reads containing >10% ambiguous bases or >50% bases with Phred quality score below 20. Reads were aligned to the human reference genome (hg38) using HISAT2 and quantified with featureCounts. Gene expression was normalized to fragment per kilobase of transcript per million mapped reads (FPKM), and Principal Component Analysis (PCA) was conducted to assess sample clustering.

Differential expression analysis was performed using DESeq2, comparing treatment conditions (25 mM glucose or 500 µM MG) to 7 mM glucose (basal control) within each cell line. Genes with |fold-change| ≥ 2 and false discovery rate (FDR) < 0.05 were considered significant. Visualization of volcano plots was performed using GraphPad Prism (Version 10.5.0) and the Venn diagram was generated using R Studio, using the VennDiagram and gridExtra packages. Pathway enrichment was conducted with QIAGEN Ingenuity Pathway Analysis (IPA; version 01-23-01).

To examine treatment effects on specific pathways, we used volcano plots to visualize DEGs. Gene sets representing glycolysis, oxidative phosphorylation, cell proliferation, DNA damage repair, and RNA processing were curated from the Kyoto Encyclopedia of Genes and Genomes (KEGG). Pathway IDs can be found in Supplemental Table 9. Genes from these pathways were plotted on the volcano plots and considered significantly if they met a |fold-change| ≥ 1.2 and an FDR < 0.05.

| | AA/B | |
|---|---|---|
| | Control (N = 83) | Case (N = 83) |
| Age (years)$^{T}$ | 62.63 ± 5.69 | 62.37 ± 7.13 |
| BMI$^{T}$ | 30.82 ± 5.72 | 28.46 ± 4.73 |
| WAA$^{T}$ | 0.78 ± 0.12 | 0.80 ± 0.11 |
| Smoking (no/yes, but quit/no)$^{x}$ | (25/36/22) | (30/29/24) |
| Alcohol (no/yes, but quit/no)$^{x}$ | (11/32/40) | (14/22/47 |
| PSA$^{T}$ | 1.17 ± 0.72 | 7.38 ± 5.01 |
| Diabetes (no/yes)$^{x}$ | (53/30) | (64/19) |
| Education | 1.86 ± 1.14 | 1.55 ± 1.11 |

| P | EA | | | P value (AA/B vs. EA) | |
|---|---|---|---|---|---|
| | Control (N = 128) | Case (N = 77) | P | Control | Case |
| 0.791 | 65.06 ± 6.20 | 63.01 ± 6.38 | 0.025 | 0.004 | 0.550 |
| 0.004 | 28.14 ± 5.34 | 29.10 ± 4.78 | 0.186 | 8.0E-04 | 0.398 |
| 0.410 | 0.04 ± 0.06 | 0.03 ± 0.02 | 0.093 | | |
| 0.523 | (59/60/9) | (37/30/10) | 0.277 | 2.0E-04 | 0.042 |
| 0.249 | (12/24/92) | (3/9/65) | 0.108 | 0.001 | 4.8E-04 |
| <0.0001 | 1.51 ± 1.14 | 4.65 ± 1.71 | <0.0001 | 0.5351 | 0.0002 |
| 0.088 | (110/18) | (67/10) | 0.994 | 3.5E-04 | 0.155 |
| 0.077 | 2.85 ± 1.24 | 2.52 ± 1.22 | 0.065 | 2.1E-08 | 5.4E-07 |

Table 2. Univariate linear regression for MG-adduct in AA/B men (n=141).

| | **Log2(CEG)** | |
|---|---|---|
| | **Coefficient (Std.Error)** | **P-value** |
| **Age** | -0.03 (0.01) | 0.001 |
| **BMI** | -0.01 (0.01) | 0.35 |
| **Prostate cancer** | | |
| No | -- | |
| Yes | 0.42 (0.12) | 0.001 |
| **Education** | | |
| High school or below | -- | |
| Above high school | -0.02 (0.13) | 0.86 |
| **Smoking** | | |
| No | -- | |
| Yes, but quit | -0.18 (0.15) | 0.27 |
| Yes | -0.26 (0.17) | 0.12 |
| **Alcohol** | | |
| No | -- | |

| | | |
|---|---|---|
| Yes, but quit | -0.02 (0.21) | 0.92 |
| Yes | -0.12 (0.19) | 0.52 |
| **Married** | | |
| No | -- | |
| Yes/lived as married | -0.02 (0.13) | 0.86 |
| **Family History** | | |
| No | -- | |
| Yes | 0.17 (0.17) | 0.32 |

Table 3. Multiple linear regression for MG-adduct in AA/B patients (n=141)

| | **Log2(CEG)** | |
|---|---|---|
| | **Coefficient (Std.Error)** | **P-value** |
| **Age** | -0.03(0.01) | 0.0003 |
| **Prostate cancer** | | |
| No | -- | |
| Yes | 0.40 | 0.001 |
| **Alcohol** | | |
| No | -- | |
| Yes, but quit | -0.005 (0.19) | 0.98 |
| Yes | -0.11 (0.18) | 0.58 |

Table 4. Univariate linear regression for MG-adduct in EA patients (n=174)

| | **Log2(CEG)** | |
|---|---|---|
| | **Coefficient (Std.Error)** | **P-value** |
| **Age** | 0.004 (0.01) | 0.55 |
| **BMI** | -0.002 (0.01) | 0.77 |
| **Prostate Cancer** | | |
| No | -- | |
| Yes | -0.01(0.10) | 0.89 |
| **Education** | | |
| High school or below | -- | |
| Above high school | 0.16 (0.10) | 0.12 |
| **Smoking** | | |
| No | -- | |
| Yes, but quit | 0.05 (0.10) | 0.61 |
| Yes | 0.02 (0.17) | 0.89 |
| **Alcohol** | | |
| No | -- | |

| | | |
|---|---|---|
| Yes, but quit | -0.06 (0.22) | 0.77 |
| Yes | 0.05 (0.19) | 0.81 |
| **Married** | | |
| No | -- | |
| Yes/lived as married | -0.03 (0.10) | 0.78 |
| **Family History** | | |
| No | -- | |
| Yes | 0.03 (0.12) | 0.83 |

Table 5. Multiple linear regression for MG-adduct in EA patients (n = 174)

| | **Log2(CEG)** | |
|---|---|---|
| | **Coefficient (Std.Error)** | **P-value** |
| **Age** | 0.004 (0.01) | 0.59 |
| **Prostate Cancer** | | |
| No | -- | |
| Yes | -0.01 (0.10) | 0.90 |
| **Alcohol** | | |
| No | -- | |
| Yes, but quit | -0.08 (0.23) | 0.73 |
| Yes | 0.03 (0.20) | 0.87 |

Table 6. Analysis of GLO1 SNPs in AA/B and EA subjects with and without cancer.

| *GLO1* SNPs | chr/pos | Bonferroni Corrected p-value Race:Tumor | FDR p-value Race:Tumor | FDR Tumor:Race | MAF AA/B-Cancer | MAF AA/B-Control | MAF EA-Cancer | MAF EA-Control |
|---|---|---|---|---|---|---|---|---|
| rs9470916 | 6:38676685 | 6.64E-01 | 1.90E-02 | 5.69E-01 | 0.03086 | 0.01205 | 0 | 0 |
| rs937662 | 6:38678593 | 8.26E+01 | 9.39E-01 | 7.77E-01 | 0.05625 | 0.06024 | 0.05147 | 0.06667 |
| rs13212218 | 6:38678633 | 7.46E+00 | 1.52E-01 | 5.69E-01 | 0 | 0 | 0.01449 | 0.004065 |
| rs3799703 | 6:38678854 | 1.13E+01 | 2.06E-01 | 6.81E-01 | 0.075 | 0.06627 | 0.07812 | 0.1218 |
| rs34971977 | 6:38680034 | 1.58E+01 | 2.68E-01 | 7.91E-01 | 0 | 0 | 0.007246 | 0.004065 |
| rs140458850 | 6:38680292 | 7.73E+00 | 1.52E-01 | 5.22E-01 | 0.01235 | 0.04217 | 0.04348 | 0.06504 |
| rs1130534 | 6:38682812 | 5.21E-05 | 2.60E-06 | 8.28E-01 | 0.07407 | 0.07229 | 0.007246 | 0 |
| rs60262339 | 6:38688674 | 4.33E-01 | 1.31E-02 | 5.22E-01 | 0.0375 | 0.01205 | 0 | 0 |
| rs1579028 | 6:38692202 | 7.65E+00 | 1.52E-01 | 7.38E-01 | 0.0125 | 0.006024 | 0 | 0 |
| rs1049346 | 6:38703061 | 4.96E-05 | 2.603E-06 | 5.97E-03 | 0.399 | 0.247 | 0.040 | 0.123 |

Table 7. Primers for GLO1 sequencing in MDA-PCA-2b and C4-2 cells.

| **Target SNP** | **Primer** | ***GLO1* Primer Sequence** | **Product Size** | **Primer Tm (°C)** |
|---|---|---|---|---|
| rs1130534 + rs4746 | Forward | 5´-GTCTAATCAGTTAAGAATAATCGACACC-3´ | 608 | 53.6 |
| rs1130534 + rs4746 | Reverse | 5´-GTGGTAAGATTTCAAAGATGTTACTTGC-3´ | | 54.9 |
| rs1049346 | Forward | 5´-CTCTTCCCATCACACTCCCG-3´ | 430 | 59.8 |
| rs1049346 | Reverse | 5´-GAGGCATAGTCCTGTGGGTG-3´ | | 59.8 |

Table 8. Sequencing results for SNPs in C4-2 and MDA-PCa-2b cells

| Cell line | SNP | Chromosome | Position | Location | Gene | Reference allele | Altered allele |
|---|---|---|---|---|---|---|---|
| C4-2 | rs1049346 | 6p21.2 | 38703061 | 5'-UTR | *GLO1* | C | - |
| C4-2 | rs4746 | 6p21.2 | 38682852 | Exon 4 | *GLO1* | A | C |
| MDA-PCa-2b | rs1049346 | 6p21.2 | 38703061 | 5'-UTR | *GLO1* | C | T |
| MDA-PCa-2b | rs4746 | 6p21.2 | 38682852 | Exon 4 | *GLO1* | A | - |

Table 9. KEGG pathways used for volcano plots

| **Physiological Process** | **Pathway** | **KEGG ID** | **Source** |
|---|---|---|---|
| Glucose Regulation | Glycolysis | hsa00010 | KEGG hsa00010 |
| Glucose Regulation | Insulin Signaling | hsa04910 | KEGG hsa04910 |
| Glucose Regulation | Adipocytokine Signaling Pathway | hsa04920 | KEGG hsa04920 |
| Glucose Regulation | Regulation of lipolysis in adipocytes | hsa04923 | KEGG hsa04923 |
| Glucose Regulation | Fatty Acid Biosynthesis | hsa00061 | KEGG hsa00061 |
| Glucose Regulation | Type 2 Diabetes | hsa04930 | KEGG hsa04930 |
| Glucose Regulation | Insulin Resistance | hsa04931 | KEGG hsa04931 |
| Oxidative Phosphorylation | Oxidative Phosphorylation | hsa03410 | KEGG hsa03410 |
| Cell Proliferation | WNT | hsa04310 | KEGG hsa04310 |
| Cell Proliferation | NFκB | hsa04064 | KEGG hsa04064 |
| Cell Proliferation | TNF | hsa04668 | KEGG hsa04668 |
| Cell Proliferation | PI3K-Akt | hsa04151 | KEGG hsa04151 |
| Cell Proliferation | MAPK | hsa04010 | KEGG hsa04010 |
| Cell Proliferation | JAK-STAT | hsa04630 | KEGG hsa04630 |
| DNA Damage | Base excision repair | hsa03410 | KEGG hsa03410 |

| | | | |
|---|---|---|---|
| DNA Damage | Nucleotide excision repair | hsa03420 | KEGG hsa03420 |
| DNA Damage | Mismatch repair | hsa03430 | KEGG hsa03430 |
| DNA Damage | Homologous recombination | hsa03440 | KEGG hsa03440 |
| DNA Damage | Non-homologous end joining | hsa03450 | KEGG hsa03450 |
| RNA Processing | RNA degradation | hsa03018 | KEGG hsa03018 |
| RNA Processing | RNA transport | hsa03013 | KEGG hsa03013 |
| RNA Processing | mRNA surveillance | hsa03015 | KEGG hsa03015 |
| RNA Processing | RNA polymerase | hsa03020 | KEGG hsa03020 |
| RNA Processing | Basal transcription factors | hsa03022 | KEGG hsa03022 |
| RNA Processing | Spliceosome | hsa03040 | KEGG hsa03040 |

Supplemental Figure 1

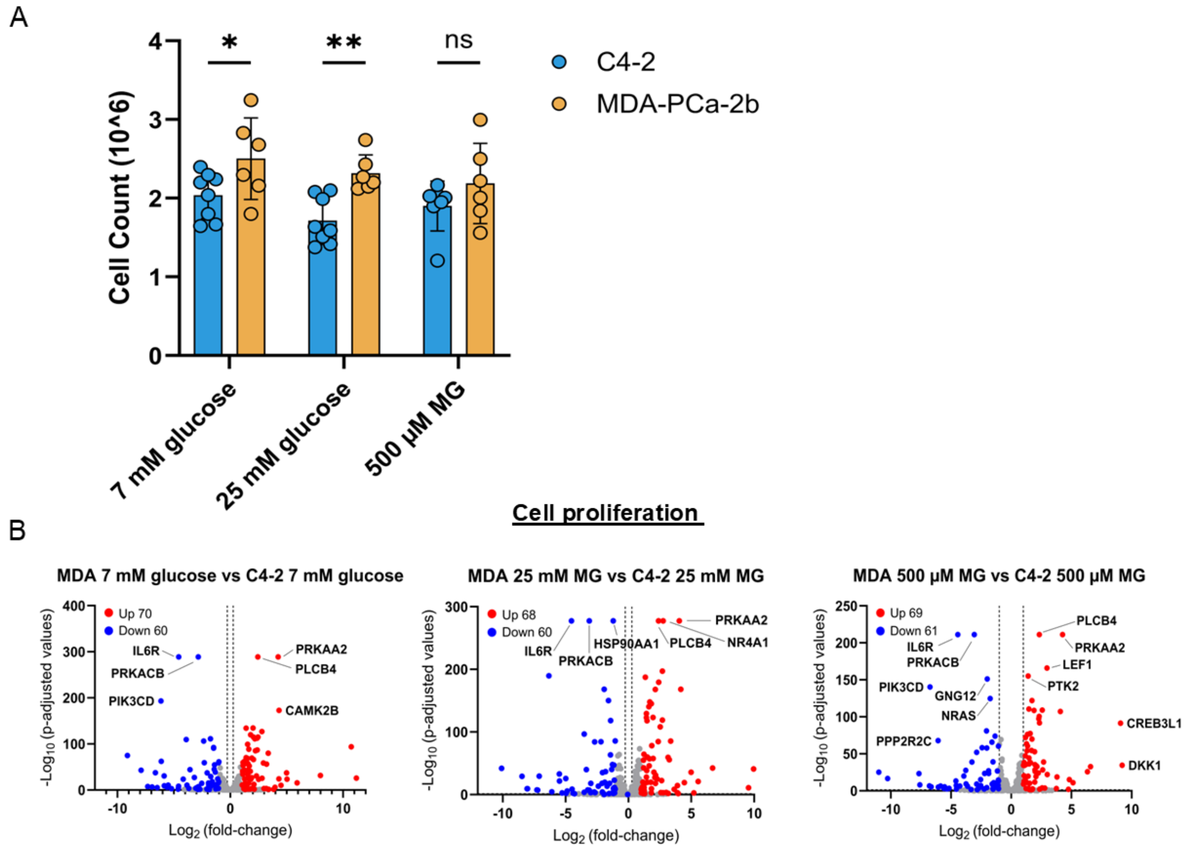


**Supplementary Figure 1. MDA-PCa-2b cells have significantly higher cell proliferation. MDA-PCa-2b cells with increased metabolic activity also have greater amounts of proliferative genes expressed, resulting in greater proliferation and expression of proliferative genes. (A)** Two-way ANOVA shows significant increase in MDA-PCa-2b proliferation in comparison to C4-2 cells after plating 1 x $10^6$ on a 10 cm dish and subjecting cells to experimental treatments within a 48-hour period. **(B)** Volcano plots were generated utilizing genes documented in corresponding KEGG pathway and showed which genes are upregulated and downregulated in hyperglycemic and high MG conditions between C4-2 and MDA-PCa-2b cell lines.
